# pmTR database: population matched (PM) germline allelic variants of T-cell receptor (*TR*) loci

**DOI:** 10.1101/2020.12.23.424097

**Authors:** Julian Dekker, Jacques J.M. van Dongen, Marcel J.T. Reinders, Indu Khatri

**Affiliations:** Department of Immunology, Leiden University Medical Center, 2333 ZA Leiden, The Netherlands.; Leiden Computational Biology Center, Leiden University Medical Center, 2333 ZC Leiden, The Netherlands.; Hogeschool, Leiden.; Delft Bioinformatics Lab, Delft University of Technology, 2628 CD Delft, The Netherlands.

**Keywords:** population, germline, allelic variants, *TR* loci, diversity

## Abstract

T-cell receptor (*TR*) germline alleles are arranged, organized and made available to the research community by the IMGT database. This state-of-the-art database, however, does not provide information regarding population specificity and allelic frequencies of the genes all four human *TR* loci (*TRA, TRB, TRG* and *TRD*). The specificity of allelic variants to different human populations can, however, be a rich source of information when studying the genetic basis of population-specific immune responses in vaccination and disease. To make *TR* germline alleles available for such population-specific studies, we meticulously identified true germline alleles enriched with complete *TR* allele sequences and their frequencies across 26 different human populations, profiled by “1,000 Genomes data”. We identified 205 *TRAV,* 249 *TRBV,* 16 *TRGV* and 5 *TRDV* germline alleles supported by at least four haplotypes (= minimum of two individuals). The diversity of germline allelic variants in the *TR* loci is highest in Africans followed by Non-African populations. A majority of the Non-African alleles are specific to the Asian populations, suggesting a diverse profile of *TR* germline alleles in different human populations. Interestingly, the alleles known in the IMGT database are frequent and common across all the superpopulations. We believe that this new set of genuine germline *TR* sequences represents a valuable new resource which we have made available through the new population-matched *TR* (pmTR) database, accessible via https://pmtrig.lumc.nl/.

## Introduction

The genomic organization of four loci of T-cell receptors (TR), i.e. alpha (*TRA*), beta (*TRB*), gamma (*TRG*) and delta (*TRD*), is complex. The four loci are distributed over three different genomic regions across two chromosomes in the human genome: *TRA* and *TRD* on chromosome 14q11.2 (with *TRD* lying within the *TRA* locus), and *TRB* and *TRG* on different arms of chromosome 7. The *TRB* and *TRD* loci are comprised of *Variable (V), Diversity* (*D*) and *Joining (J)* genes, whereas *TRA* and *TRG* loci contain *V* and *J* genes only. The polypeptides, encoded by functionally rearranged *TRA* and *TRB* loci, combine to form a TRαβ receptor, whereas functionally rearranged *TRD* and *TRG* loci form the TRγδ receptor, both containng an antigen-recognition domain. The recombination of *V(D)J* genes has the potential to generate many millions of TR molecules having a unique antigen binding specificity [1]. The *V(D)J* recombination process is directed by “recombination signal sequences” (RSS), short highly conserved DNA stretches, present at each recombination site of the *TR* genes, i.e. downstream to *V*, upstream to *J*, and at both sites of *D* [2,3].

*TR* genes harbor inter-individual germline allelic variants, causing different individuals to be able to produce different receptors. As these different allelic variants are shared within confined human populations [4], they contribute also to more extreme diversity of receptors at the population level [5]. These population-specific germline variations have been shown to introduce varying disease prevalences in specific population [6–9]. For example, in Asian and Caucasian populations, *TRBV17* plays a pivotal role in Influenza A virus specific T-cell immunity [10]. Consequently, to understand (population-specific) immune responses, a catalogue of population-wide observed TCRs is crucial. Till date, there is, however, only one database that reports all the alleles for the *TR* locus: the International ImMunoGeneTics information system (IMGT) [11,12]. But, this database does not report allelic frequencies or population statistics and, moreover, reported alleles are mostly profiled from Caucasian populations [13,14].

To enrich the catalogue of TR germline genes with population information, we relied on the “1,000 Genomes (G1K)” dataset (https://www.internationalgenome.org/), derived from cell samples of 2,548 individuals across five different ethnicities. We are not the first in doing so. Yu *et al* created the Lym1k database for immunoglobulin (*IG*) and *TR* loci, also from the G1K data using their AlleleMiner tool [14]. They, however, did not provide any information on the reliability of the (newly) identified alleles as was the link to population information not retained. Moreover, not all relevant components of each *TR* locus were stored, i.e. they neglected the *D, J, C* genes and the RSSs. Also, they were not able to profile all *TR* genes as they used a previous version of the G1K dataset (i.e. a mapping to GRCh37 being liftover to GRCh38).

Here, we identified the alleles for all components of all four *TR* loci, i.e. the *V, D, J, C* genes as well as the RSSs, report reliability scores for the differently detected alleles as well as population information of each allele, and present an online accessible database containing this information which we called the “population-matched germline allelic variants of T-cell receptor loci” database; or in short, the *pmTR* database. To realize this, we have developed an automated pipeline to profile all the *TR* alleles from the G1K data. The pipeline returns the sequences of alleles, frequency of alleles, as well as the population distribution of each allele among 26 different populations profiled in G1K resource. The resulting alleles are manually curated and made available via GitHub and the online database (www.pmTRIG.com), including population information and confidence levels to provide access for the community. We have also enabled a BLAST search on the database to directly use our germline alleles in further research.

## Results

### Population matched germline *TR* alleles (pmTR) database application

We identified population-specific alleles in all four *TR* loci (*TRA*, *TRB, TRG* and *TRD*) using pmAlleleFinder pipeline (Methods), where each allele is supported by at least four haplotypes for 2,548 individuals belonging to 26 populations representing five continents (**Table S1**). All the alleles identified from G1K are identified to create a population matched *TR* (pmTR) database. We identified two to three times more new alleles then present in the standard reference IMGT database for all the genes in the variable genes of all *TRA* and *TRB* loci (**Table 1**). These alleles were divided into three allele sets (AS1, AS2, or AS3) based on different confidence levels (Methods). AS1 are alleles already present in the IMGT database; AS2 are novel alleles that are frequent in the populations (supported by at least 19 haplotypes); and AS3 are novel but rare alleles (supported by 4 to 18 haplotypes). The pmTR database further contains meta information about the alleles such as the support of haplotypes for each (sub)population (**Tables S2, S3, S4, and S5)**. We have also identified the Recombination signal sequences (RSS) for each gene and the corresponding variants for each allele. The heptamers and nonamers in the RSS sequence turned out not to be conserved for most *TR* genes resulting in lower recombinant frequencies for those genes [22]. Similar to *IG* pseudogenes [13], we also identified conserved heptamers for *TR* pseudogenes suggesting the role of RSS in recombination of pseudogenes in T-cell repertoire [23] (**Table S6**).

**Table 1:** Number of alleles in different functional gene segments in *IG* loci. AS1 (Known), AS2 (Frequent) and AS3 (Rare) are major confidence levels.

|  | AS1 | AS2 | AS3 | Total | IMGT |
| --- | --- | --- | --- | --- | --- |
| *TRAV* | 54 | 75 | 76 | 205 | 87 |
| *TRAJ* | 64 | 18 | 17 | 99 | 71 |
| *TRAC* | 1 | 2 | 1 | 4 | 1 |
| *TRBV* | 68 | 82 | 99 | 249 | 118 |
| *TRBD* | 1 | 0 | 0 | 1 | 3 |
| *TRBJ* | 16 | 1 | 0 | 17 | 16 |
| *TRBC* | 1 | 2 | 3 | 6 | 4 |
| *TRGV* | 13 | 30 | 23 | 66 | 16 |
| *TRGJ* | 5 | 2 | 1 | 8 | 6 |
| *TRGC* | 13 | 8 | 7 | 28 | 13 |
| *TRDV* | 5 | 5 | 2 | 12 | 5 |
| *TRDD* | 3 | 0 | 0 | 3 | 3 |
| *TRDJ* | 4 | 2 | 0 | 6 | 4 |
| *TRDC* | 1 | 0 | 2 | 3 | 1 |

### Variable alleles in the IMGT database are partial and are frequently present in all ethnicities

Mapping the pmTR alleles to the known alleles is instrumental in this setting for assessing the frequency and the population specificity of the known alleles as such information will be helpful in understanding the population-specific response to vaccines and disease. We found that 60-100% of the IMGT alleles for functional variable genes in each *TR* loci were identified in the pmTR database. The functional *V* gene alleles in the IMGT database are not complete as we found 42 of 87 *TRAV* and 53 of 118 *TRBV* alleles to be partial i.e. they do not comprise a leader sequence. In most cases only the first, or first two, alleles of the *TR* genes are sequenced completely. Moreover, the majority of the *TR* genes in the IMGT had only one allele recorded, implying that a complete *TR* germline allele resource is not available for the research community. When mapping pmTR and IMGT alleles only over the *V* exon region, an increase in the known alleles (AS1 category) was observed. Moreover, looking at the super-population distribution of the IMGT alleles in our pmTR database, we found that a majority of these alleles (>90%) are shared among all the superpopulations (**Figure 1**). Moreover, more than 90% of the mapped IMGT alleles are supported with at least 100 haplotypes and are frequently present in all the superpopulations (**Figure 3, Table S2-S5**).

**Figure 1:**
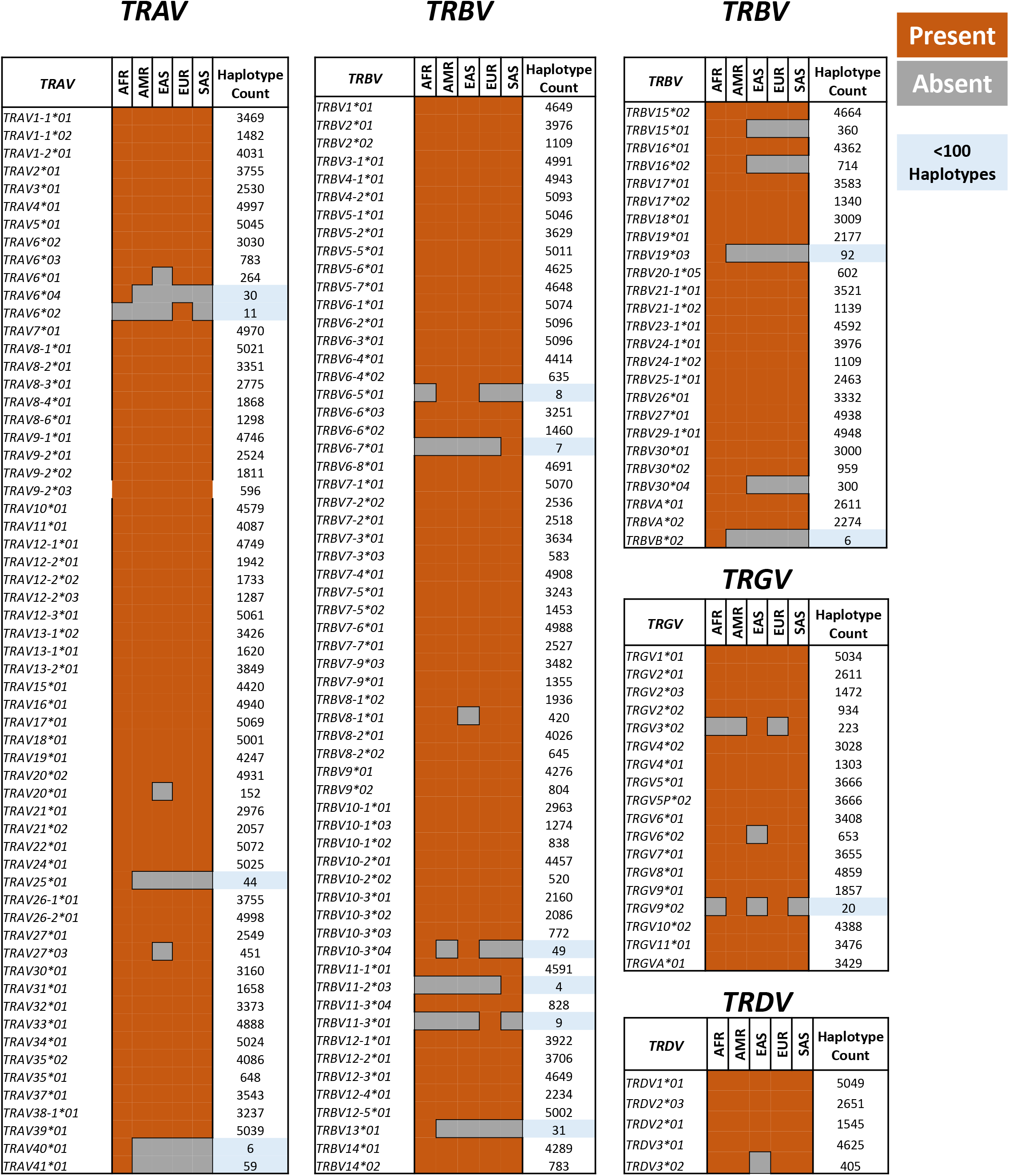
Superpopulation distribution of the known (IMGT) *V* gene alleles in all four *TR* loci. The IMGT alleles identified in the pmTR database are shown here with the IMGT identifiers. The presence or absence of the populations specificity for these alleles are shown in the orange and grey shade respectively. The last column shows the frequency of these alleles. The blue shaded frequencies depict the alleles with <100 haplotypes. These alleles are frequently present in all the superpopulations.

### The majority of the novel *V* gene alleles are one mutation away from known alleles

To gain information on the number of new mutations added by the novel *V* gene alleles to the mutating positions in the known germline alleles, it was important to estimate the differences that are added by the mutations in the Leader region and *V* exon separately as only *V* exon rearranges with (*D)J* genes to generate T-cell repertoire. For complete *V* genes (Leader + *V* exon), we found that 81% (113/139) of the novel *TRAV* alleles and 88% (142/161) of the novel *TRBV* alleles are one mutation away from their known alleles (**Table 2**), i.e. they only have one different mutation with respect to a known IMGT allele. These mutated positions are randomly distributed over the *V* gene and do not show specific patterns.

**Table 2:** Number of alleles with count of new mutating positions as compared to the existing databases. Complete sequence includes leader sequence and the *V* region for all the *V* genes/ This is important to realize that 42 *TRAV* and 53 *TRBV* alleles are partial and do not contain the leader sequence.

| Mutating positions | TRAV |  | TRBV |  |
| --- | --- | --- | --- | --- |
|  | Complete sequence | *V* region only | Complete sequence | *V* region only |
| 0 | 66 | 71 | 88 | 104 |
| 1 | 113 | 114 | 142 | 132 |
| 2 | 22 | 17 | 14 | 10 |
| 3 | 3 | 2 | 5 | 3 |
| 4 | 1 | 1 | 0 | 0 |

### Novel alleles are unique to super-populations

Opposite to the known alleles (AS1 category), which are shared between all the population, novel alleles for *TR* loci (AS2 and AS3 categories) are unique to specific superpopulations. Very few novel *TR* alleles are shared among all the superpopulations. One-third of the novel alleles in all the four *TR* loci constitute of African-specific alleles (**Figure 2**). As shown in Figure 1, only 3 of 66 *TRAV* African-specific alleles are known in the IMGT database. Two-third of the novel African specific alleles are rare, i.e. supported by less than 19 haplotypes. In line with this, none of the 34 population-specific *TRAJ* alleles are known in the IMGT database (**Figure 2A**). However, we do not observe a similar pattern for the *TRBD, TRBJ* and *TRBC* alleles (**Figure 2B**). *TRBV,* on the other hand, follows a similar pattern as the *TRAV* alleles; half of the novel alleles are African-specific of which a majority is rarely present in Africans (**Figure 2B**). Moreover, half of the ‘AFR shared’ *TRAV* and *TRBV* alleles are common between African and American superpopulations (**Figure 2A**), suggesting a role of intermixing due to migratory events.

**Figure 2:**
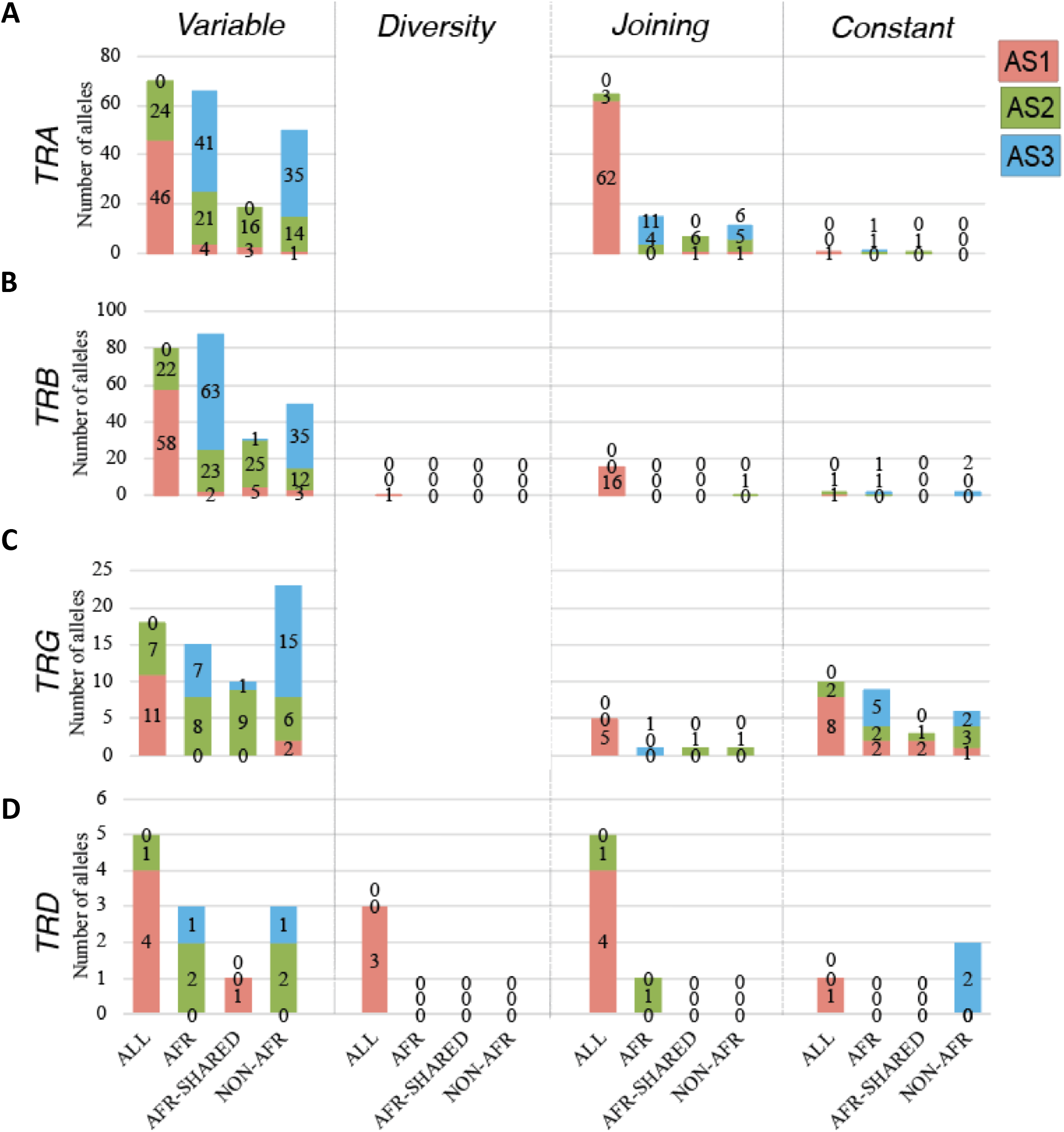
Superpopulation distribution of the alleles for variable, diversity, joining and constant genes in A) TRA; B) TRB; C) TRG; and D) TRD loci. The population plots are represented for all superpopulations (ALL), Africans only (AFR), Africans shared with one of the other superpopulations (AFR Shared) and ‘Non-AFR where alleles are present in one of the populations other than Africans. Red label background indicates AS1 alleles, green AS2 and blue AS3 alleles.

*TRG* and *TRD,* being the smallest *TR* loci, have fewer novel alleles as compared to the *TRA* and *TRB* loci. However, novel *TRGV* alleles often belong to Non-African populations (**Figure 2C**). The *TRG* locus has the highest number alleles for constant genes across all four *TR* loci, a majority of which belongs to the rare category (AS3) (**Figure 2C**). The *TRD* locus is the most conserved locus as very few novel alleles were found for the *TRDV, TRDJ* and *TRDC* genes (**Figure 2D**). Summarizing, we find a similar superpopulation distribution across all four *TR* loci despite their size difference and varying levels of duplication.

### A majority of Non-African alleles are specific to Asian populations

The superpopulation distribution of novel *TRA*, *TRB* and *TRG* alleles show that these are specific to the Non-African populations (**Figure 2**). In fact, a majority of them belongs to the East and South Asian populations (**Figure 3**). Interestingly, these alleles are not specific to any of the Asian populations. Very few alleles were shared between Asian and American-European populations, suggesting an exclusive nature of the *TR* loci in these populations. The larger number of Asian-specific alleles suggests an exceptional diversity in Asia which went unnoticed in the current databases.

**Figure 3:**
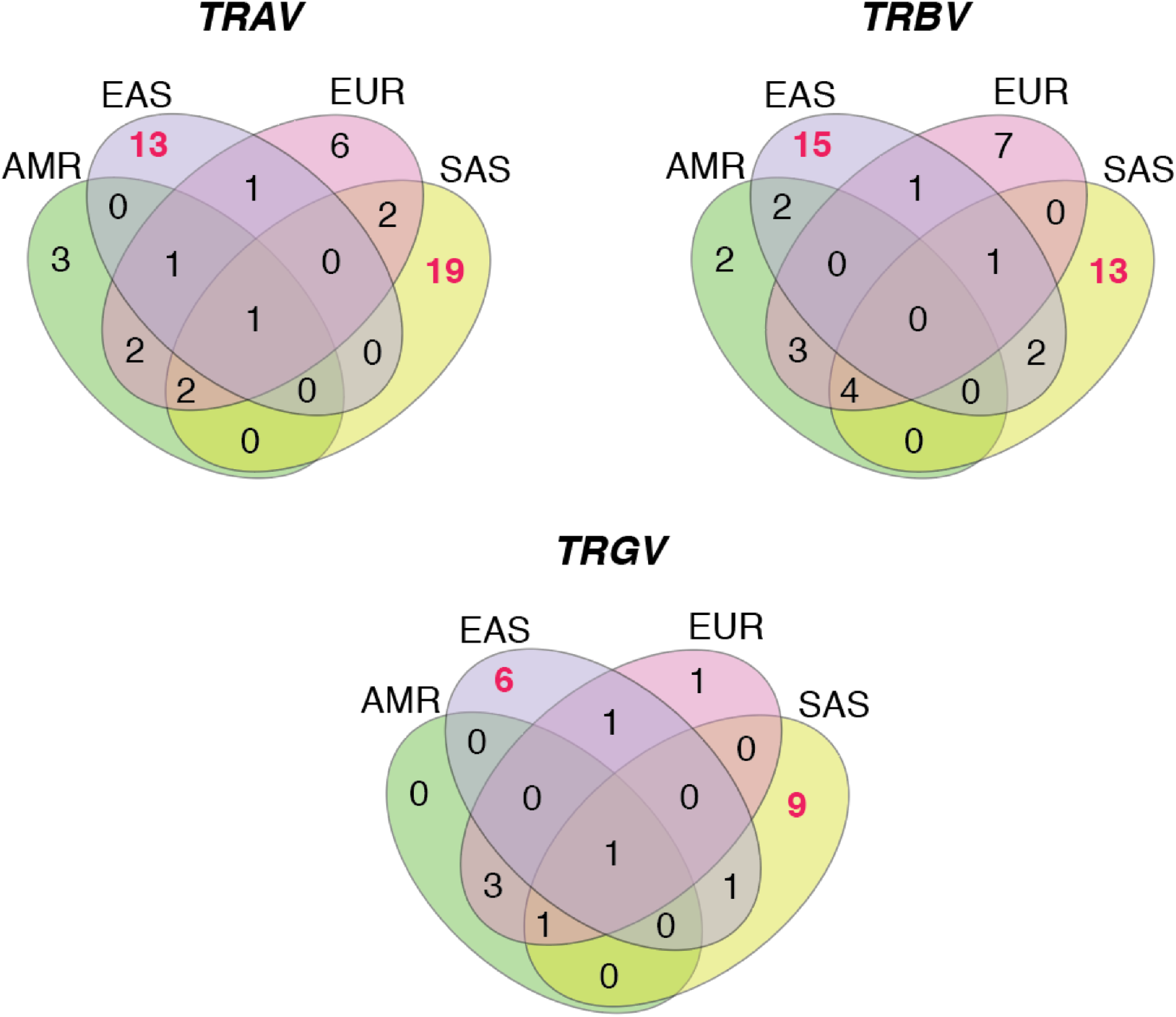
Superpopulation specificity of the Non-African alleles in A) TRA; B) TRBV; and C) TRG genes. The genes specific to East and South Asian populations are shown in red color.

### Genomic organization of the *TR* loci governs the evolutionary dynamics

A principle component analysis (PCA) was performed on the entire span of all four *TR* loci to visualize the genetic diversity among different loci. We found that the Africans are highly diverse in all the four loci, whereas other superpopulations are comparatively similar to each other (**Figure 3**). Interestingly *TRA* and *TRB* follow similar patterns despite being on different chromosomes. On the other hand, in the *TRG* and *TRD* loci, we observed multiple clusters of individuals from all the superpopulations having unique variability as compared to the variation in the individuals belonging to African populations. Despite that the *TRD* locus lies within the *TRA* locus, it does not follow a similar variability as the *TRA* locus, implying that selection pressure has been different when comparing the *TRA* and *TRB* loci with the *TRG* and *TRD* loci. This may have been governed by the size of loci and functional aspects of the TCRαβ vs. TCRγδ molecules.

To investigate the population structure in more detail, we calculated the population differentiation for each *TR* loci separately. We found a comparatively different structure between the loci as compared to the genetic diversity assessed using PCA in **Figure 4A**. We found Africans to be the most diverse in the *TRB* and *TRG* loci whereas Americans and Europeans are ancestor clades for the *TRA* and *TRD* loci. The population structure is in accordance with the chromosomal organization of these loci, unlike the genetic diversity visualized in the PCA plot. Interestingly, in all the *TR* loci, an early separation of Mexicans (MXL) and Peruvians (PEL) populations is observed (**Figure 4B**). Similar to the population structure in *TRG* loci, the relatedness of the PEL population was also observed in the *IG* loci [13].

**Figure 4:**
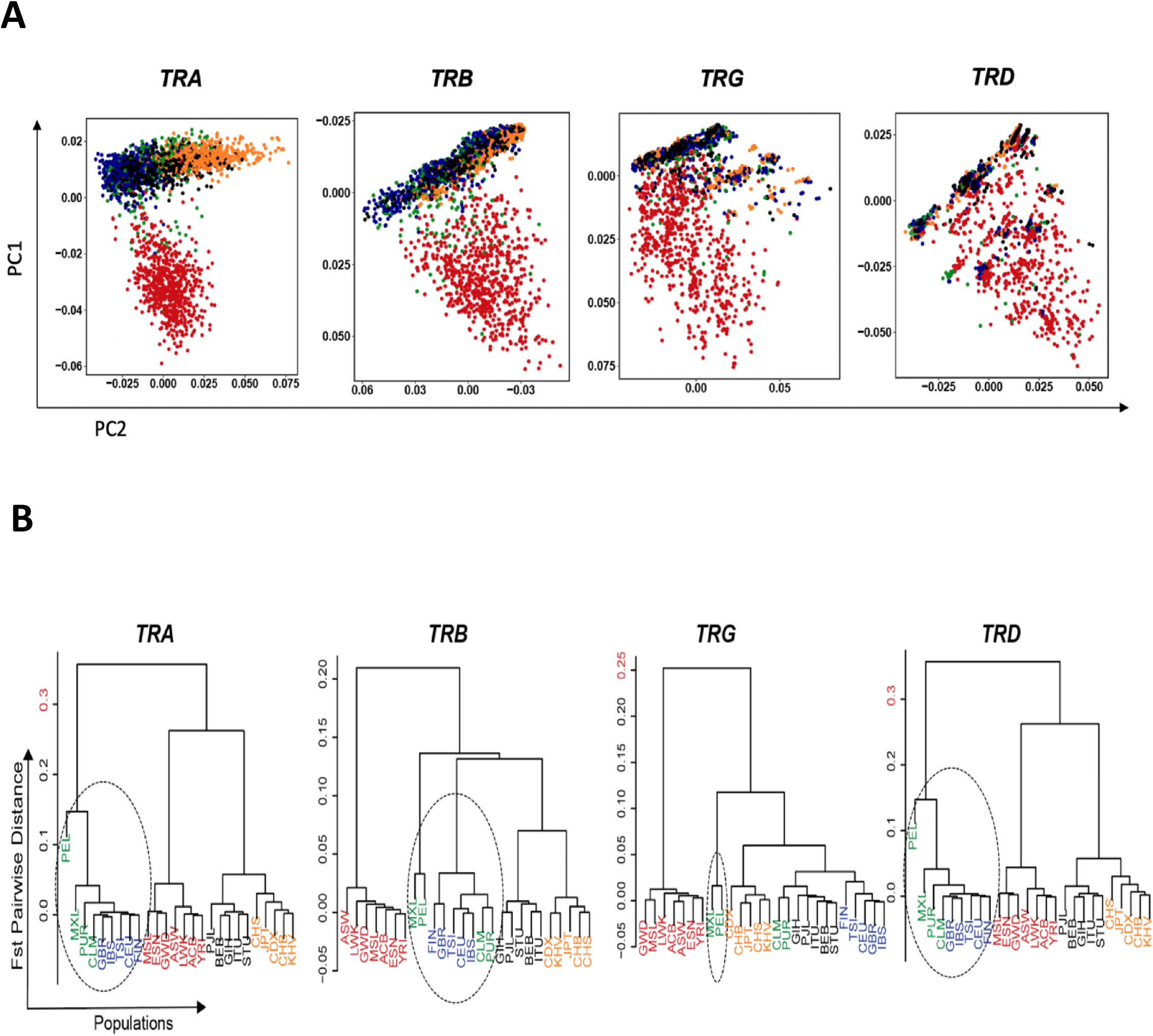
Genetic diversity and population structure in five superpopulations for *TR* loci. **A) Separate PCA plot of all four *TR* loci based on the polymorphisms in the complete locus.** Each dot represents a sample and each sample is colored based on the superpopulation they belong. **B) Pairwise population distribution calculated by F_ST_ Matrix is represented as a cladogram for each locus namely *TRA, TRB, TRG* and *TRD.*** Five superpopulations are colored as Africans in red; Americans in green; East Asians in orange; Europeans in blue and South Asians in black.

## Discussion

We performed a comprehensive analysis of *TR* germline alleles identified from 2,548 individuals available in the 1,000 Genomes data belonging to 26 populations across five different ethnicities. The *TR* germline alleles from the 1,000 Genomes data were also identified previously by Yu et al and compiled in the Lym1K resource. This resource, however, does not provide information on the allelic frequencies and population-specificities of the alleles [14]. In addition, we not only used a minimum haplotype count (i.e. 4 haplotypes) to identify alleles using our automated pipeline “pmAlleleFinder”, but also categorized the pmTR alleles in three confidence levels. Moreover, potential false-positive germline alleles in operationally indistinguishable (OI) genes were also eliminated by manually assessing the mutating positions between the alleles belonging to OI genes [13]. Finally, the filtered alleles for *TR* loci were compiled in the pmTR database (hosted via https://pmtrig.lumc.nl/). Applying these rules and checks have provided us with genuine TR germline alleles.

The assignment of the alleles to three confidence categories provided us with a notion of accuracy for the pmTR alleles. 60-100% of the IMGT alleles for *Variable* genes were also recorded in the pmTR database. The coverage even increased to >80% of the IMGT alleles when the mapping was performed across the *V* exon region only. A complete overlap between the pmTR and IMGT *Variable* gene alleles is hampered by the number of individuals with which the pmTR database is created, i.e. 2,548, as compared to thousands of individuals profiled by the IMGT database over last decade. However, in the pmTR database, all genes/alleles from one individual are profiled, whereas the IMGT database has been compiled with few (rearranged) genes/alleles of many different individuals. We found that >90% of the IMGT alleles are shared between all five superpopulations, with very few alleles specific to one or more of the superpopulations, suggesting that the IMGT database lacks rare population-specific alleles. Having only alleles that are shared uniformly and frequently across all superpopulations poses limitations for immune-response studies that are aimed at understanding the genetic basis of difference between and among different human populations [4,15].

Most of the novel pmTR alleles were identified in African populations. >90% of these novel alleles are not captured by the IMGT database. In contrast to IG alleles [13], where African-Shared alleles were the second largest group of alleles [13], the Non-African alleles are the second largest group for TR alleles. A majority of these Non-African alleles are specific to the Asian populations, suggesting a divergence of *TR* loci, resulting in unique alleles across different ethnicities. We also observed a similar trend in the evolutionary dynamics of the independent *TR* loci represented by the genetic diversity in the PCA plots. The African individuals showed the highest genetic diversity in all the loci. However, we found that the genomic organization governs the population structures of the four *TR* loci. *TRA* and *TRD* showed similar patterns with the Americans and Europeans being the most diverse, whereas the African populations are the most variable for the *TRB* and *TRG* loci. Here it should be noted that the *TRD* and *TRG* loci are the smallest loci, and, hence, the genetic diversity and the population structure can be affected by the size of the loci.

Taken together, we meticulously reported curated germline alleles across 5 ethnicities containing 26 subpopulations, resulting in ~150% more alleles as compared to the known alleles within the IMGT database. This enriched resource can be used for the repertoire studies to understand (population-specific) immune response dynamics.

## Star Methods

### Data source

The 1,000 Genomes data (G1K) (March, 2019 release; http://www.1000genomes.org; GRCh38 assembly) in the form of phased variant cell format (VCF) was used for retrieving the *TR* germline alleles. Phased variants for GRCh38 (a recent release for the 1,000 genomes) were used as they were the most recent and comprised almost all the genes of the *TR* locus as compared to the GRCh37 assembly. The full release of the data set was collected from 2,548 cell samples from diverse ethnic groups that have a uniform distribution of individuals across populations. The samples are classified in five super populations i.e. Africa, America, East Asia, Europe and South Asia, that are further subdivided into 26 populations (7 African, 4 American, 5 East Asian, 5 European and 5 South Asian populations) with a minimum of 61 and a maximum of 113 samples per population (**Table S1**). The phased VCF format of the data comprises information of both parental (forward) and maternal (reverse) chromosomes for each sample.

### Terminology and Nomenclature

With the term *“haplotype’’”* we refer to a gene present on one strand (inherited from a single parent) in one individual. Therefore, there are two haplotypes, one on the positive and one on the negative strand, with exactly the same or different polymorphisms. *“Allele”* refers to a haplotype from multiple individuals consisting of the same variants across the complete gene sequence. *“Mutations”* are genetic mutations that occurred to form different alleles (also denoted as allelic variants) of the same gene. The IMGT nomenclature is used to name genes, and this name is extended with a numbering to refer to the different alleles, for example the 01 and 02 alleles of the *TRAV1* gene are referred to as *TRAV1_01, TRAV1_02.* IMGT alleles are denoted with an asterix, such as *TRAV1*01*, *TRAV1*02*. The alleles were sorted in descending order such that the first allele is supported by the maximum number of haplotypes.

### An automated pipeline to identify germline alleles from G1K data

The pmAlleleFinder pipeline is used to infer the alleles of the genes of interest i.e. *V*, *D*, *J* and *C* genes and RSSs from the input VCF file. The pipeline results in a list of alleles for each gene separately along with the population information of each allele in a separate file (**Figure S1**). It finds all possible haplotypes for each gene, merges them into alleles and counts the haplotype frequency of each such allele. The alleles can be filtered by the user through defining a minimum number of haplotype support. The pipeline is developed in python and R with an additional possibility to automatically identify if pmTR alleles are present in the IMGT database. The pipeline is not limited to the identification of alleles for *TR* genes only and, given a phased vcf file, can be used to find population-based alleles for any gene.

### Allele confidence levels

The alleles obtained from the G1K resource were classified into three major confidence levels (allele set (AS) 1-3):

*AS1 (known):* G1K alleles with a minimum support of 4 haplotypes and identified in the IMGT databases. This AS1 allele set has the highest level of confidence as the alleles are observed in the G1K resource as well as in the IMGT database.

*AS2 (frequent novel alleles):* G1K alleles with a minimum support of 19 haplotype (at least ten individuals). These alleles represent a set of newly identified alleles that are frequent.

*AS3 (rare novel alleles):* G1K alleles that have a haplotype support between 4 and 18 (supported by two to nine individuals). Despite the rarity of these alleles, we believe them to be genuine as the chance that 4 *identical* haplotypes within 5,096 independent haplotypes are caused by sequencing errors is highly unlikely.

As few *V* genes are paralogous [15,16], the mapping of short reads to such genes can be erroneous, influencing the subsequently derived alleles. Called mutations on the alleles of such genes can thus easily be false positives, even after using stringent parameters. Therefore, we denote these genes as *operationally indistinguishable* (OI) genes. As these genes can be recognized based on their sequence similarity [17], we generated a neighbor-joining (NJ) tree for all *V* genes on the *TRA*, *TRB, TRG* and *TRD* loci, separately. The genes sharing a clade with a short branch length, i.e. 0.02, are called OI genes (**Figure S2**); and the corresponding alleles OI alleles.

### Filtering false positive alleles

The G1K alleles were scrutinized manually: 1) alleles with stop codons were removed from the final set, and 2) alleles within OI genes where removed when they had a mutation that is shared with alleles for the other OI genes as it points towards a mis-alignment of a read (when the mutation is present in the IMGT databases across multiple alleles it is not filtered).

### Population annotation of alleles

G1K alleles are annotated with superpopulation information (**Tables S2-S5**) into four categories: 1) ALL, present in all superpopulations; 2) AFR, only present in Africans; 3) AFR SHARED, present in African and at least one of the other superpopulations, but not all; and 4) NON-AFR, present in at least one of the superpopulations but not in Africans.

### pmTR online database

The online pmTR database front-end is made with ReactJS in combination with the Neo4j graph database back-end to load and display all genes [18,19], respective alleles, population frequencies and confidence levels (AS1-AS3). Genes can be searched on the basis of their name or nucleotide sequence enabled by a BLAST search [20] (**Figure S3**). The pmTR is hosted on https://pmtrig.lumc.nl/.

### Genetic diversity and migration events

The VCF file of the complete individual locus, i.e. *TRA* (Chr14 [21621904, 22552132), *TRB* (Chr7 [142299011, 142813287]), *TRG* (Chr7 [38240024, 38368055]) and *TRD* (Chr14 [22422546, 22466577]) was obtained. The SNPs from the coding and non-coding region from each locus were independently subjected to a principal component analysis (PCA) using the R Bioconductor package ‘SNPRelate’ [21]. The pairwise population differentiation, quantified by the fixation index (F_ST_), was calculated based on levels of differentiation in polymorphism frequencies across populations. F_ST_ is proportional to the evolutionary branch length between each pair of populations. A Neighbor joining tree was used to visualize the F_ST_ distances between populations.

## Acknowledgements

We acknowledge 1,000 Genomes project for making the data publicly available. As we have used a specific locus of the chromosomes 7 and 14, we have complied with the 1,000 Genomes policies for the publication of data.

## Authors contributions statement

The study was conceptualized by JJMvD. The pipeline was developed by IK which was automated by JD. JD performed the data acquisition, data analysis and organization and development of the database. IK and MJTR supervised the analysis. All the authors wrote the manuscript and designed the figures. All the authors approved the final version of the manuscript.

## Availability of data

The source code of the automated tool to identify population-matched alleles and automated mapping to known resources is available via GitHub (https://github.com/JulianDekker/PMalleleFinder). The database is hosted via a website at (www.pmTRIG.com).

## Supplementary Figure legend

**Figure S1:**
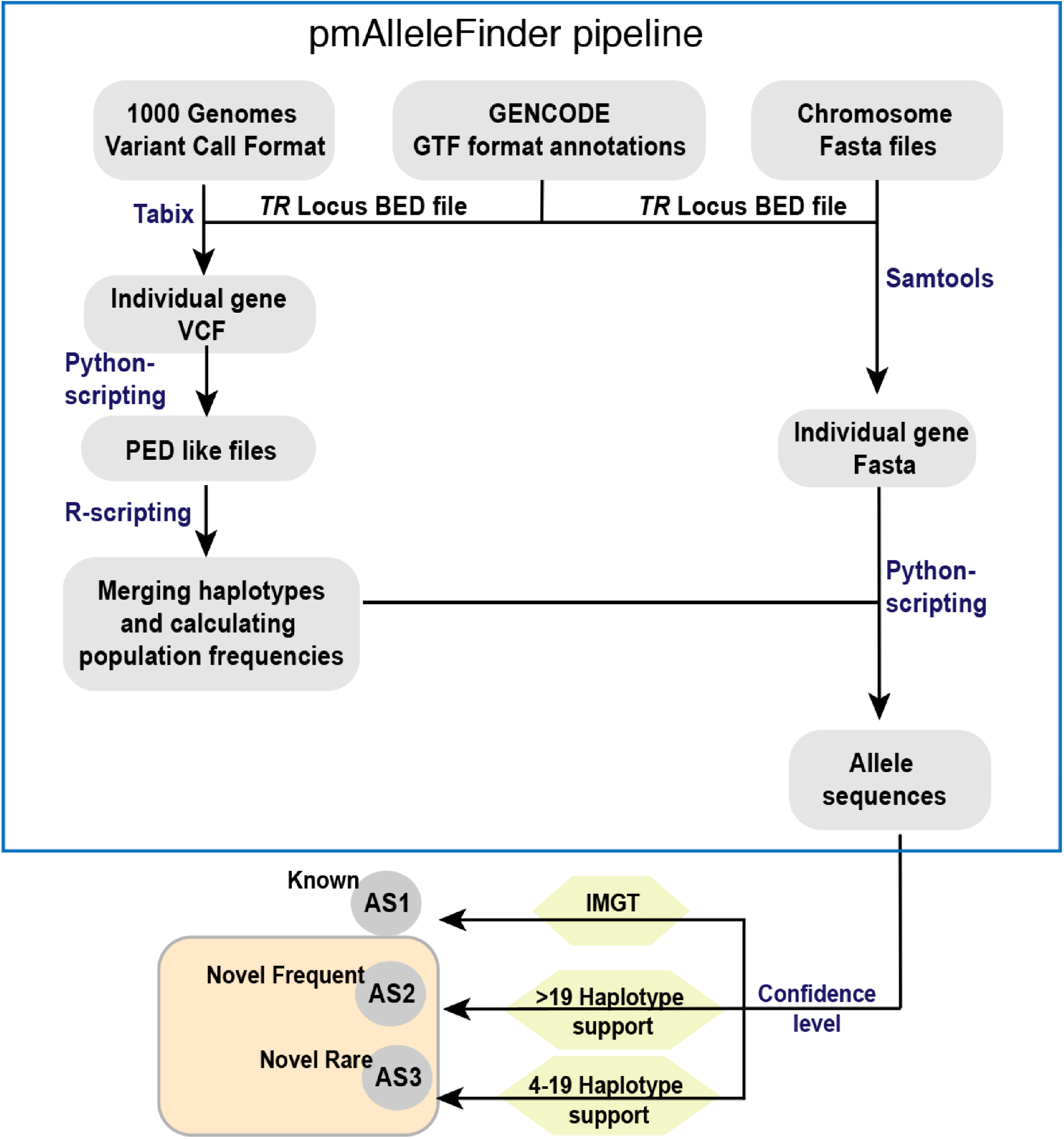
pmAlleleFinder: an automated pipeline to identify population matched alleles. The workflow depicts the VCF, GENCODE GTF format and Chromosome Fasta files are needed as input. The combination of python and R scripts are to identify haplotypes, population frequency and the allele sequences. These sequences are further divided into three categories using automated pipeline developed in Python.

**Figure S2:**
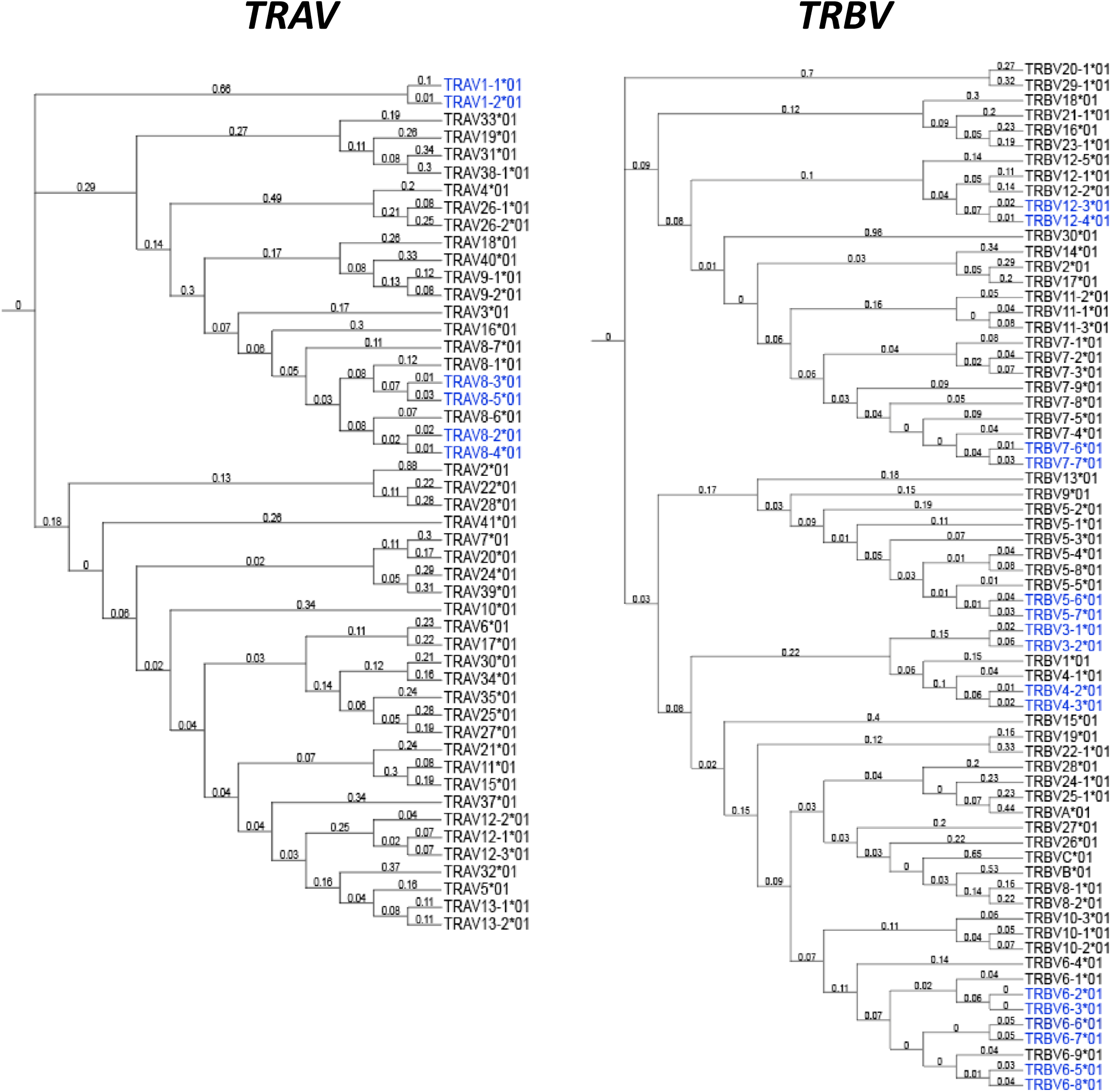
Neighbor Joining tree for the *TRAV* and *TRBV* genes from IMGT database to identify duplicated genes. The genes separated by a small distance are considered as operationally indistinguishing genes that are marked in blue in the tree. The alleles from these genes are scrutinized manually for shared mutating positions.

**Figure S3:**
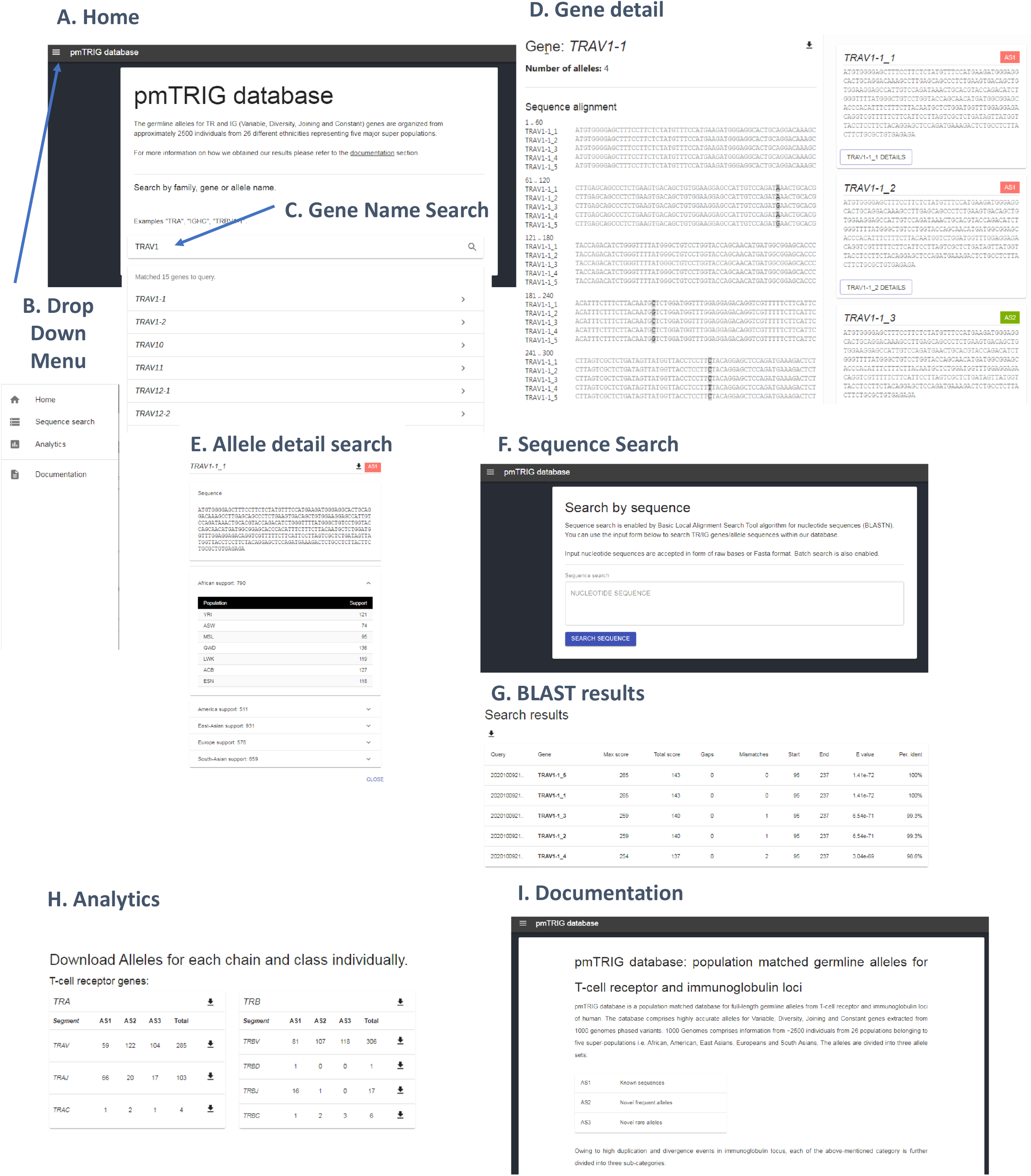
pmTRIG database: The database with organized population matched germline alleles, identified from G1K samples, for the *TR* and *IG* loci. All the different components of the website are shown i.e. **A)** Home screen; **B)** Drop-down Menu; **C)** Box with gene name search; **D)** result for a gene search; **E)** Information provided for each alleles; **F**) Sequence search screen; **G)** Output of sequence search in BLAST format; **H)** Analytics page providing downloads of genes for each confidence category for each loci; and **I)** Documentation page for the database.

